# A Refined Open State of the Glycine Receptor Obtained Via Molecular Dynamics Simulations

**DOI:** 10.1101/668830

**Authors:** Marc A. Dämgen, Philip C. Biggin

## Abstract

Fast neurotransmission is mediated by pentameric ligand-gated ion channels. Glycine receptors are chloride-selective members of this receptor family that mediate inhibitory synaptic transmission and are implicated in neurological disorders including autism and hyperekplexia. They have been structurally characterized by both X-ray crystallography and cryo electron microscopy studies, with the latter giving rise to what was proposed as a possible open state. However, recent work has questioned the physiological relevance of this open state structure, since it rapidly collapses in molecular dynamics simulations. Here, we show that the collapse can be avoided by a careful equilibration protocol that reconciles the more problematic regions of the original electron-density map and gives a stable open state that shows frequent selective chloride permeation. The protocol developed in this work provides a means to refine open-like structures of the whole pentameric ligand-gated ion channel superfamily and reconciles the previous issues with the cryo-EM structure.

## Introduction

The superfamily of pentameric ligand-gated ion channels (pLGICs) are key players in the communication between nerve cells. Situated in the postsynaptic membrane they mediate fast synaptic transmission. As such, they are fundamental for cognitive processes and are involved in a range of neurological disorders such as Parkinson’s or Alzheimer’s disease which makes them important drug targets^1-3^. An atomistically-detailed understanding of the functional states of these receptors would be extremely valuable for structure-based drug design.

In the resting state, the ion-conducting pore within the transmembrane domain (TMD) is closed. Upon the binding of neurotransmitters to their extracellular domain (ECD), the receptors undergo conformational changes that lead to the opening of the central pore^4^, thereby allowing ions to pass through, leading to a change in the membrane potential and thus translating the chemical neurotransmitter signal into an electrical nerve signal. From this ion-conducting active state, they then undergo a conformational change to a non-conducting desensitized state with the neurotransmitter still bound ^5^. All members of the pLGIC superfamily share a very similar overall architecture^6,7^ with cation-selective channels acting as excitatory and anion-selective channels as inhibitory receptors.

The glycine receptor (GlyR) is an inhibitory member of this superfamily that is selective for chloride ions and the endogenous agonist is the amino acid glycine. Glycinergic inhibition is localized primarily in the brain stem and spinal cord where it is essential for motor coordination and pain processing^8-10^. Consequently, GlyRs are a promising drug target for spasticity, inflammatory pain and hyperekplexia (startle disease)^11-13^. GlyRs have been the focus of both functional^14-16^ and structure studies^6^ and indeed various structures of the glycine receptors with agonist, antagonist and modulators bound have been resolved with cryo-electron microscopy (cryo-EM)^17^ as well as X-ray crystallography^18-20^ and have been annotated to closed, open and desensitized states.

From these structures and those of other members of the superfamily alongside a large amount of mutagenesis work (see Thompson *et al*.^21^ and references therein), it became apparent that there are two main constriction points in the channel pore, namely a ring of five hydrophobic residues (usually leucine) at the 9’ position of the pore-lining M2 helices and in the case of the GlyR, a ring of five proline residues at the -2’ position. The so called L9’ gate, located in the middle of the TMD closes the channel in the antagonist (strychnine) bound structures (PDB codes: 5CFB and 3JAD). The P-2’ gate is near the intracellular mouth of the channel pore and gates the channel in the desensitized-like structures (5TIN, 5TIO, 5VDH, 5VDI and 3JAF). Among these structures, the glycine bound cryo-EM structure 3JAE is of particular interest due to its unusually wide-open pore conformation relative to other open-like structures of the pLGIC superfamily, particularly at the -2’ position. The structure 4HFI^22^ of the proton-activated bacterial *Gloeobacter violaceus* (GLIC) at pH=4 and the 3RIF structure^23^ of the invertebrate glutamate-gated chloride channel (GluCl) from *Caenorhabditis elegans* were classified as open-like pLGIC structures. The distance from the pore centre to the backbone C*α* at the -2’ position is about 2Å smaller for these two structures than for the wide open 3JAE structure^24^.

MD simulations have provided useful insight into the functional annotation of states to ion channel structures, which is difficult to infer from structural information alone. An important concept in this context is hydrophobic gating^25-27^. Hydrophobic gating occurs in narrow hydrophobic pores where the energetically favoured expulsion of water prohibits ion permeation. This means that a pore whose dimensions are theoretically large enough to allow for ion permeation is not necessarily ion-permeant due to hydrophobic effects. Pore hydration has been suggested as a proxy for ion-permeability^28^ and used to annotate ion channels as functionally open or closed.

Trik *et al*.^28^ and Gonzalez-Gutierrez *et al*.^24^ independently annotated the wide open 3JAE GlyR structure as open. Furthermore, Gonzalez-Gutierrez et al. found the 3JAE wide open pore to have a four times higher single channel conductance than the experimental value (although it should be noted that the simulations were performed at 37 °C, while experiments were conducted at room temperature). Moreover, blocking the channel pore with picrotoxin in their simulations still resulted in ion conduction whereas no currents were observed experimentally. This substantiates the notion that the wide open 3JAE atomic model has an overly dilated pore. In these simulations, artificial restraints were applied to the protein backbone, such that it cannot move freely and stays very close to the original cryo-EM model. Interestingly, if no such restraints are applied, the structure undergoes a so called hydrophobic collapse to a distinct state with a significantly smaller pore radius^29^. This behaviour has also been observed in simulations of other open-like structures of the pLGIC family^30-32^. They all have in common that the hydrophobic collapse is initiated via the non-polar interaction of the 9’ gate residues.

We hypothesized that the wide open atomic model 3JAE that has been fitted into the cryo-EM density map may not correspond to an energetically stable representation of the open state under physiological conditions. We took the view that other models that agree with the cryo-EM density map may exist and these may be conformationally stable. To find alternative open state structures, we developed a careful equilibration protocol where we give the protein maximum freedom to explore other open state configurations while preventing the pore from collapsing. We then release all restraints on the pore and show that the structure found through this careful equilibration procedure stays open with a hydrated pore that allows selectively for frequent chloride ion permeation. The model provides alternative conformations of the leucine side-chains that are not discernible in the original electron density map. Moreover, we provide a structural explanation as to why the structure found in our MD-simulations is a stable open state, whereas the wide-open structure modelled into the cryo-EM density map undergoes a hydrophobic collapse as observed by us and many others. The results reconcile previous apparently conflicting points of view and provide a way forward to model transitions between states in the pLGIC family.

## Results

### The MD-refined structure is physically open, hydrated and selectively permeant to chloride ions

We designed a careful equilibration protocol comprised of a stepwise reduction in restraint forces, followed by a series of flat-bottomed harmonic cross-channel restraints designed to keep the pore open (see **Methods**). The flat bottom potential only exerts a force when the Cα atoms of opposite pore-lining residues come closer to each other than in their distance in the open-state cryo-EM structure 3JAE^17^. We applied this protocol to our model of the human α1 GlyR (see **methods**). These restraints were maintained for 150 ns and the final frame was then used to seed three-independent production simulations of 300 ns duration. All simulations gave stable open-pore configurations, did not exhibit hydrophobic collapse and exhibited similar pore profiles.

Comparison of the pore radius profiles of our MD structures with the wide open cryo-EM structure (3JAE) shows the mean radius of the MD structures is slightly narrower at the P-2’ ring, but would still allow for the permeation of hydrated chloride ions (**Fig. 1a, Supplementary Fig. 1 and Supplementary Movie 1**). The MD structures are thus clearly not in a desensitized state with the P-2’ constriction being 2 Å wider than the currently highest resolution crystal structure of a desensitized state (PDB code:5TIN^19^). Examination of the pore region of a representative simulation structure (the structure with minimal C*α* RMSD to the average structure) coloured by the hydrophobicity of the amino acids surrounding the pore (**Fig 1b**) revealed that the hydrophobicity was highest in the L9’ region, which is considered the site of a hydrophobic gate in the resting state of the receptor^33,34^. Since previous work has suggested that a radius of less than ca. 4-5 Å in hydrophobic regions can lead to local dehydration and thus be a prohibitive energetic obstacle for ion permeation^35^ we assessed the hydration and ion permeation events.

**Fig. 1.**
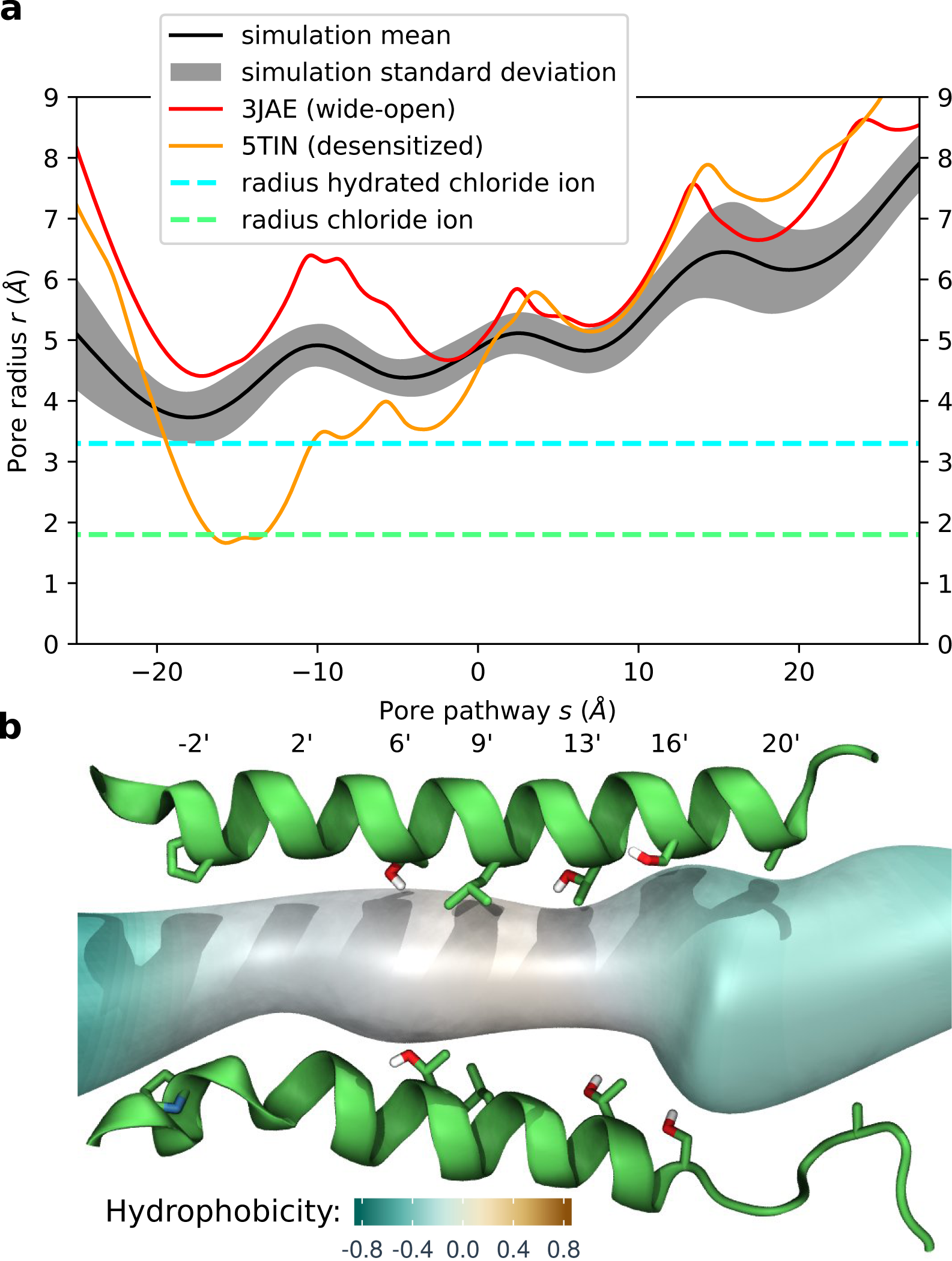
Pore profile of GlyR structures **a** A typical pore radius profile obtained from a production run (mean – black, standard deviation - grey) alongside with the profile of the cryo-EM wide open structure 3JAE (red) as well as the desensitized crystal structure with currently highest resolution 5TIN (orange). The radii of a dehydrated and a hydrated chloride ion are indicated by dashed green and cyan lines, respectively. The pore is positioned in this and subsequent figures such that the L9’ ring is located at s=0. The GlyR structure from the simulation is physically open and theoretically allows permeation of hydrated chloride ions. **b** Corresponding pore region of a representative structure from the simulation (minimal Cα RMSD to the average structure). M2 helices with pore lining residues (only 2 opposed subunits shown) together with the pore volume coloured by the hydrophobicity of the surrounding based on the Wimley-White hydrophobicity scale rescaled from -1 (very hydrophilic) to 1 (very hydrophobic).

The ion densities obtained over 300 ns of unrestrained simulation (**Fig. 2a**) reveal that chloride ions can indeed penetrate the transmembrane pore and show selectivity over sodium ions, which preferentially occupy regions in the upper part of the extracellular domain. To resolve the underlying dynamics over time, we analysed the z-coordinates of water, chloride and sodium ions inside the pore within 5 Å of the ion channel axis (**Fig. 2b**). Chloride ions frequently penetrate the transmembrane pore region, while sodium only rarely accesses this region. Over the 300ns we count 52 permeation events for chloride and a single permeation event for sodium. In two further repeats 53 and 49 chloride along with 1 and 0 sodium permeations are observed over 300 ns, respectively (see **Supplementary Fig. 2**). This is in line with the experimental estimation of a ratio of 28:1 selectivity of chloride over sodium^36^. Furthermore, the whole channel pore is hydrated throughout the simulation (as well as in the other two repeats), demonstrating that the observed pore radius at the hydrophobic L9’ gate is above the threshold forhydrophobic gating.

**Fig. 2.**
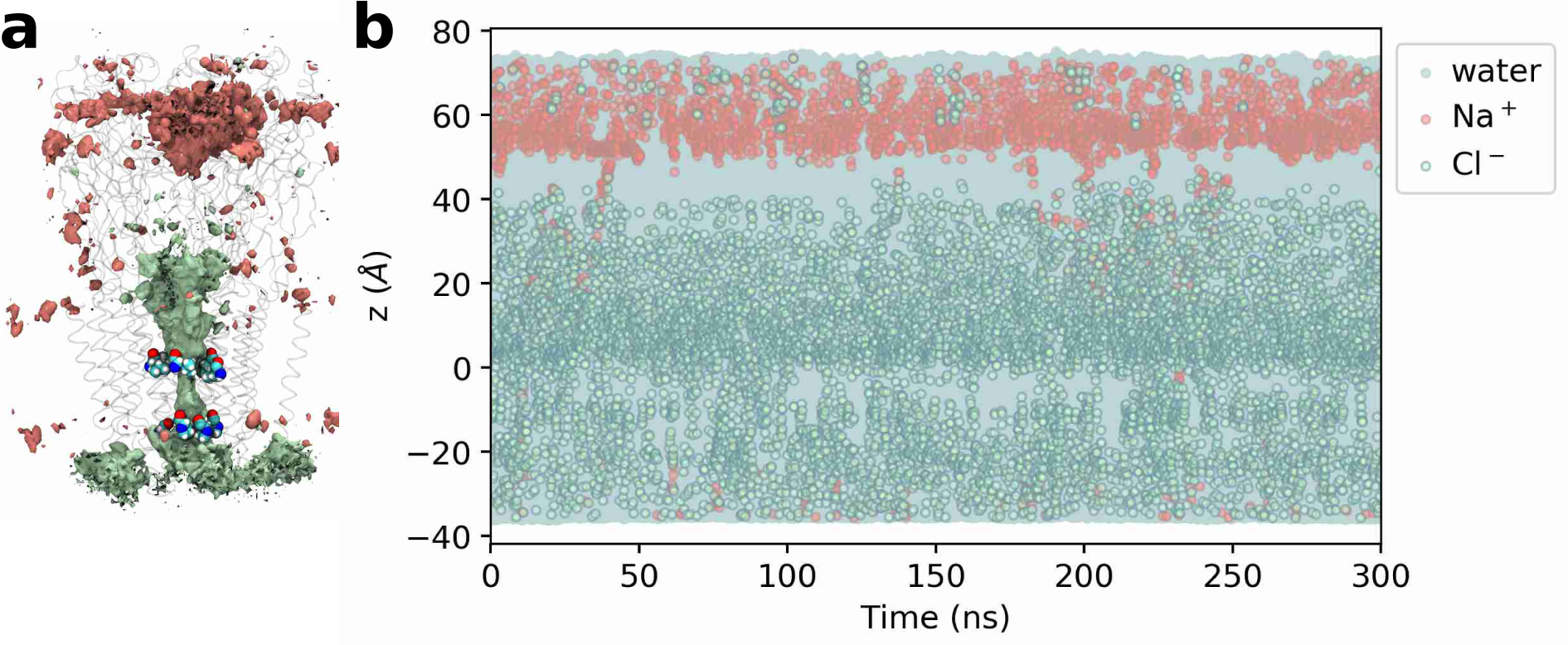
Ion densities and trajectories of water and ions within 5 Å of the channel axis. **a** Densities of chloride (green) and sodium (red) ions over a 300 ns production run. The isosurfaces shown represent a density value of 0.5 particles/nm3. Transparent ribbons indicate the receptor backbone. The L9’ ring and P-2’ ring residues are shown explicitly in van der Waals representation. The densities show chloride occupancy in the transmembrane pore and moreover, show selectivity for chloride over sodium in this region. **b** Trajectories of water as well as chloride and sodium ion z-coordinates within 5 Å of the channel axis inside the pore over 300 ns (represented by blue, green and red circles, respectively). The 0 point of the z-axis is positioned at the L9’ ring and the receptor structure in (a) is aligned and scaled correspondingly to allow for spatial orientation along the z-axis. The whole channel pore is wetted throughout the simulation (dewetted regions would appear as white stretches). While chloride frequently penetrates into the transmembrane pore, sodium does not, again demonstrating the chloride selectivity for this channel.

### The open conformation is stabilized by the leucine gate residues filling a hydrophobic void at the subunit interface

Having annotated the receptor state as functionally open, the question that arises is: Why does the structure maintain a conformationally stable open state? The key conformational changes that occur during the equilibration phase with pore restraints are depicted in **Fig. 3**. Taking a cross section at the level of the L9’ gate reveals that at the beginning of the equilibration phase, there are substantial voids between the subunit interfaces near the leucine residues (**Fig. 3a**). During the equilibration phase, the leucine residues that originally point slightly more towards the pore centre, rotate further away from the central axis and bury themselves deeply into these hydrophobic voids and thus lock the subunit interfaces, as well as the neighbouring M1 helix, strongly together (**Fig. 3b and Supplementary Movie 2**). With L9’ = L261 being located on the M2 helix of the principal (+) subunit, the corresponding hydrophobic void is formed by I257, V260 and T264 belonging to the M2 helix of the principal (+) subunit, L255, T258, T259, T262 belonging to the M2 helix of the complementary (-) subunit as well as L233, I234 and L237 of the M1 helix of the complementary (-) subunit. Across the pLGIC superfamily, all the corresponding positions are occupied by hydrophobic residues (**Supplementary Fig. 3**) and thus the hydrophobic pocket is functionally preserved. This suggests that the open state of the whole superfamily is characterized by the 9’ hydrophobic residues (L or I/V in some cases) being buried in these hydrophobic voids.

**Fig. 3.**
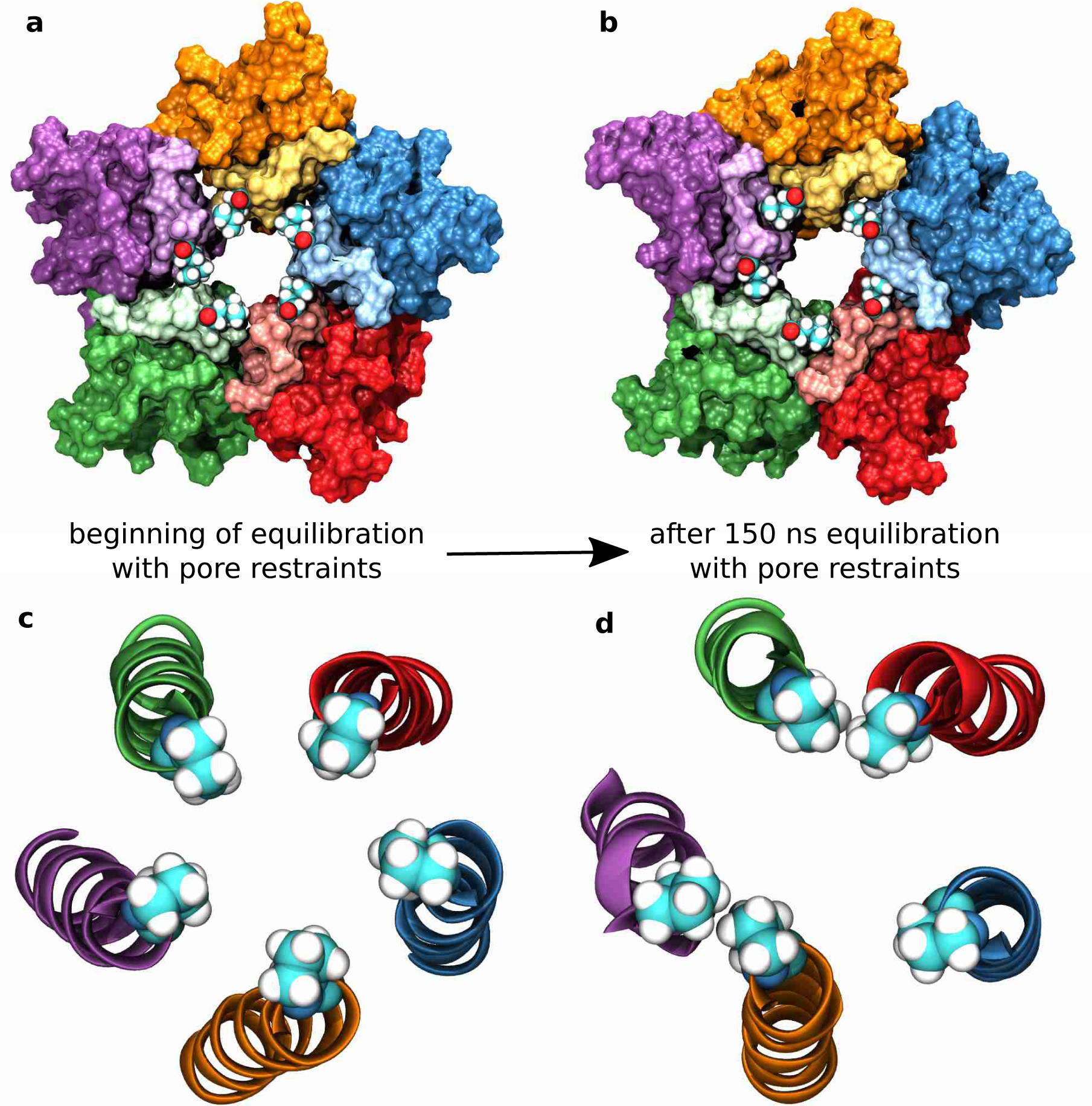
Critical changes in pore geometry. **a** and **b** View from the extracellular towards the intracellular matrix. The five subunits are coloured differently with residues that normally form the hydrophobic binding pocket around 2’ Leu in brighter colours. While at the beginning **a** the five hydrophobic voids at the subunit interfaces are empty and the L9’ residues are orientated slightly more towards the pore centre. **b** After 150 ns of equilibration with pore restraints the 9’ residues have buried themselves deeply into these hydrophobic pockets and make close contact with the surrounding residues in this pocket, thus firmly locking the structure into an open state conformation. **c** and **d** View of the M2 helices with P-2’ gate shown explicitly from the intracellular towards the extracellular side. While at the beginning, the structure is five-fold symmetrical, after 150 ns of equilibration with pore restraints, an asymmetric arrangement has been attained.

We also note that the interaction between the L9’ residue and the methyl-group of the T6’ residue of the complementary subunit leads to the T6’ hydroxyl group facing towards the centre pore. This makes the profile more hydrophilic at the 6’ ring and thus facilitates pore hydration and ion permeation. Thus, we postulate that the (+) L9’ – (-) T6’ hydrophobic interaction plays an important role in the gating mechanism of GlyR. The T6’ position is highly conserved across anion pLGICs (**see Supplementary Fig. 3**). For anionic pLGIC members, the (+) L/I9’ – (-) T6’ interaction could thus be an additional way to fine-tune pore hydration in the gating process.

While the L9’ residues lock the pore in an open conformation, the P-2’ gate residues rearrange from a symmetric conformation (**Fig. 3c**), which is enforced by the five-fold symmetry restraints when constructing the cryo-EM model, to an asymmetrical conformation (**Fig. 3d**). In this arrangement, two vicinal proline residues form hydrophobic interactions with each other, increasing the distances toward the non-interacting vicinal proline residues, such that the overall asymmetric conformation withstands a hydrophobic collapse.

### The cryo-EM based structure collapses due to a sub-optimal orientation of the leucine gate residues

Having shown that the key to a stable open state structure is the occupation of all five hydrophobic voids at the subunit interface by the L9’ gate residues, it is easy to explain why the original cryo-EM conformation undergoes a hydrophobic collapse. In the original cryo-EM conformation (see **Fig. 3a**), the L9’ residues are not buried in the hydrophobic voids at the subunit interface, but rather point slightly more towards the pore centre. In simulations that start from this conformation, the presence of explicit water together with protein dynamic flexibility at a physiological temperature then renders these hydrophobic voids extremely unstable. Collapse usually occurs when two subunits that form a void come closer together and push the corresponding leucine residue towards the pore centre (**Supplementary Movie 3**). It is also possible that a leucine residue randomly ends up in a hydrophobic void. However, the overall collapse is a chaotic process, such that at least two (but usually more) hydrophobic voids collapse with the corresponding leucine residues being pushed towards the pore centre, decreasing the pore radius significantly (**Fig. 4a**). The pore lining M2 helices come closer to each other, such that the P-2’ interactions increase, leading to a further collapse of the P-2’ ring (**Fig. 4b**).

**Fig. 4.**
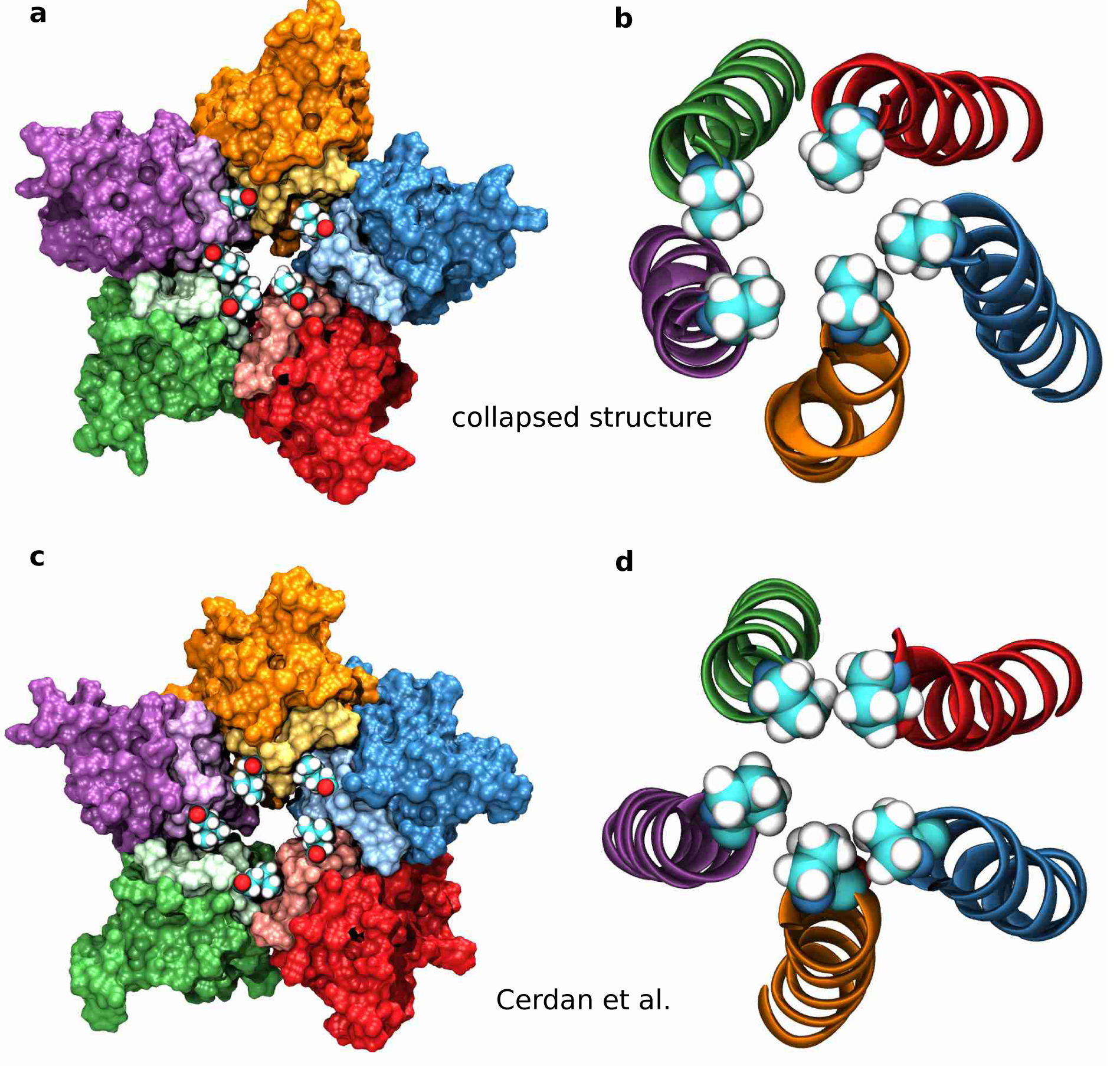
Structural features of the collapsed state. **a** and **b** Representative structure from a collapsed simulation. **a** While two hydrophobic pockets end up accommodating the corresponding leucine residues, the other three hydrophobic voids undergo a hydrophobic collapse, where the two subunits come closer together and push the leucine residues towards the pore centre and decrease the pore radius significantly. **b** With the pore lining M2 helices coming closer together, the P-2’ residues undergo a hydrophobic collapse due to their strong hydrophobic interactions. **c** and **d** Representative structure from a simulation by Cerdan et al. **c** Two hydrophobic pockets fail to accommodate their corresponding L9’ residue, leading to a much smaller pore radius. **d** Without all L9’ gate residues locking the structure into a stable open state, the P-2’ residues come so close to each other that their strong hydrophobic interactions lead to a hydrophobic collapse in the P-2’ ring region as well.

A recent report^29^ presented an ion-permeable state of the glycine receptor obtained from MD simulations. We argue that this state merely corresponds to a collapsed structure. On a structural level, it shows the key features of a collapsed state (see **Fig. 4c and 4d**). At the L9’ gate, two hydrophobic voids are collapsed and the P-2’ ring is also collapsed. The pore radius profile resembles that of our collapsed simulation with the L9’ gate pore radius being slightly larger, but the P-2’ gate pore radius slightly smaller. The authors reported a single chloride ion permeation event over a total unrestrained simulation time of 500 ns from two repeats. Chloride permeation, however, is not impossible for a collapsed state. **Fig. 5c** illustrates that occasionally chloride ions can penetrate the transmembrane pore region. For the simulation with a collapsing pore presented here, we count 3 chloride ion permeation events over 300 ns. Nevertheless, this is more than an order of magnitude less than the 52 chloride permeation events over 300 ns observed in our stable open state simulation presented here. The chloride ion density obtained from our collapsed simulation has a value below 0.5 particles/nm^3^ between the L9’ and the P-2’ gate. Hence, no green isosurface is visible in this region in **Fig. 5b**. In two further simulation repeats undergoing a hydrophobic collapse (see **Supplementary Fig. 4 and Supplementary Fig. 5**), we observe 3 and 0 chloride permeations over 300 ns unrestrained simulation time, respectively. Considering these results, we conclude that the structure reported by Cerdan *et al*. does not correspond to a stable open state, but rather to a collapsed state.

**Fig. 5.**
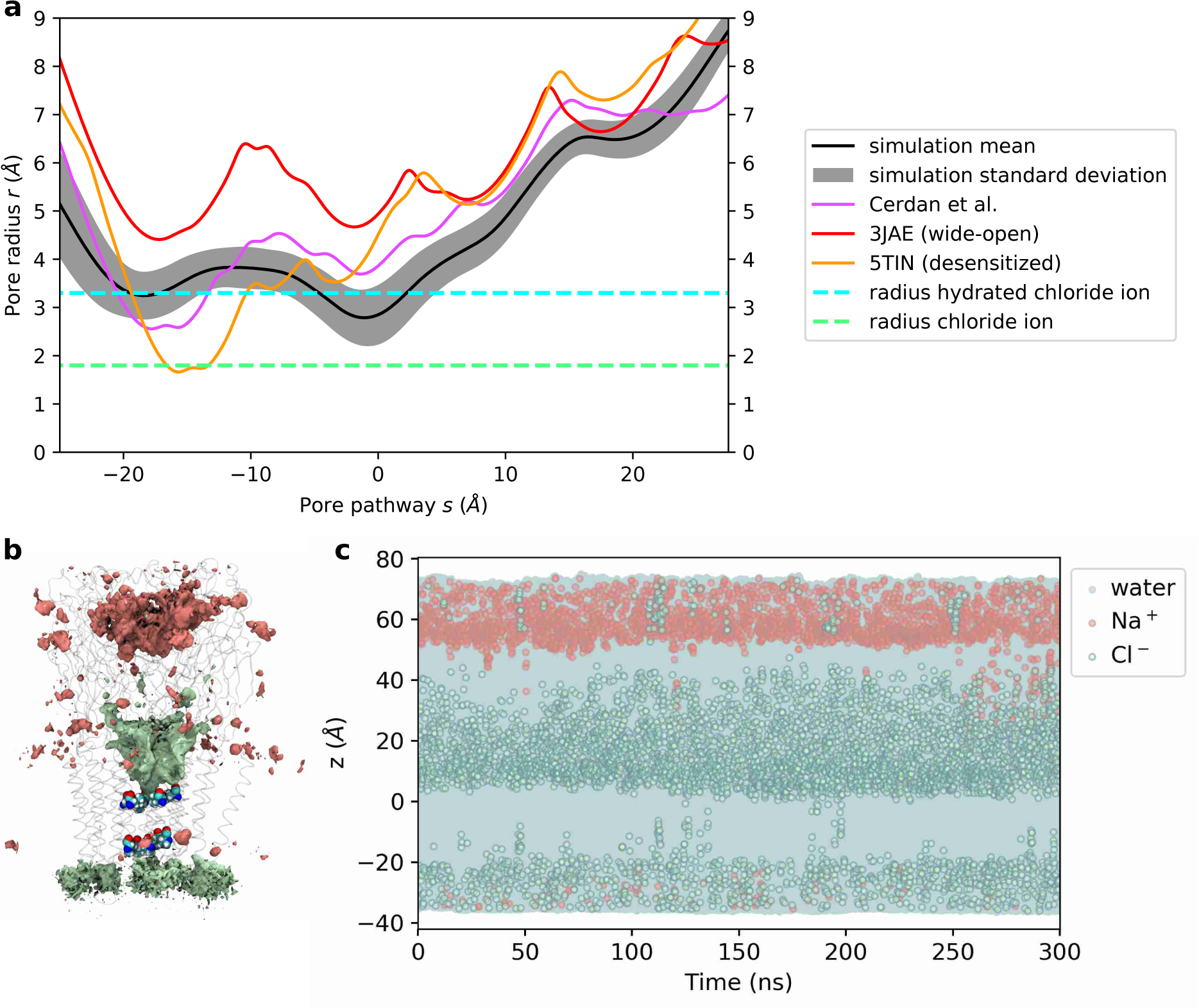
Pore profile and ion permeation of a hydrophobic collapsed state. **a** Pore radius profile of a hydrophobic collapse simulation (mean – black, standard deviation - grey) alongside with the profile an MD-structure obtained by Cerdan et al. (purple) as well as the cryo-EM wide open structure 3JAE (red) and the desensitized crystal structure with currently highest resolution 5TIN (orange). The radii of a dehydrated and a hydrated chloride ion are indicated by dashed green and cyan lines, respectively. **b** Densities of chloride (green) and sodium (red) ions over 300 ns production run. The isosurfaces shown represent a density value of 0.5 particles/nm^3^. Transparent ribbons indicate the receptor backbone, The -L9’ ring and P-2’ ring residues are shown explicitly in van der Waals representation. **c** Trajectories of water as well as chloride and sodium ion z-coordinates within 5 Å of channel axis inside the pore over 300 ns (represented by blue, green and red circles, respectively). The receptor structure in a is aligned and scaled correspondingly to allow for spatial orientation along the z-axis. The whole channel pore is wetted throughout the simulation (dewetted regions would appear as white stretches), but chloride ions access the transmembrane pore much less than in the stable open state simulation.

## Discussion

Simulations of open-like states of the pLGIC superfamily undergo a hydrophobic collapse once artificial restraints on the protein structure are released^29-32^. This is why it has become common practice to apply restraints on the protein backbone to artificially fix its position^24,28,37^. Recently,^38^ reported that while simulations starting from one structure of the 5-HT_3_ receptor (I2, PDB: 6HIQ) gave the familiar hydrophobic collapse, simulations starting from another (F, PDB: 6HIN) allowed ions and water to flow throughout the trajectory and that wetting was correlated to rotation of L9’ out of the pore lumen. The 6HIN structure shows that the L9’ leucines are indeed rotated away from the pore axis (**Supplementary Fig. 6**), consistent with our refined model of the GlyR and supporting the view that this corresponds to an open state. By forming strong hydrophobic interactions with residues of the M2 and M1 transmembrane helices of the complementary subunit, the leucines at L9’ lock the pore into a stable open conformation.

It is insightful in this context to revisit the original electron density map of the wide open 3JAE structure from cryo-EM^17^ paying particular attention to the areas of the key conformational changes that stabilize the open state, namely the L9’ and P-2’ region. Close inspection shows no density for the orientation of the L9’ gate side chains (**Supplementary Fig. 7a**), meaning that their atomic coordinates are purely a result of computational modelling and that alternative leucine side-chain conformations are equally probable.

Inspection of the density at the P-2’ gate region, also shows that the map is of particularly poor resolution in this region (**Supplementary Fig. 7c and 7d**) and likely corresponds to the high mobility of this region^17^. The data would be consistent with an ensemble of asymmetric P-2’ gate conformations which, when averaged, result in a symmetric distribution of electron density. Consistent with this, our simulations confirm a higher mobility in this area, as measured by the root mean square fluctuation (RMSF) of C*α* atoms (**Supplementary Fig. 8**). While typical RMSF values for C*α* atoms in *α*-helices are around 0.7 Å, these values are significantly increased near the P-2’ gate (> 1 Å). This means more conformational diversity of the intracellular pore mouth is accessible at a physiological temperature of 37 °C. Furthermore, the density map in the P-2’ ring region shows that there is density for proline further towards the pore centre (**Supplementary Fig. 7b**), suggesting that on average the pore radius might in reality be ca. 1 Å or so smaller than the unusual wide open –2’ Pro radius of the modelled atomic structure. This is an interesting discrepancy between the density map and the fitted model, as our simulations also suggest a smaller P-2’ pore radius of comparable magnitude (**see Fig. 1a**). Moreover, the real space correlation between the modelled structure for 3JAE and the density map is particularly meagre in this region, with R0’ peaking with a very low correlation below 0.4 (**Supplementary Fig 7e**). This means that the atomic model 3JAE is of poor quality in the M1-M2 linker and the lower M2 region, where the P-2’ gate is located.

In conclusion, the results presented here demonstrate the potential of MD simulations to refine structures of channel proteins derived from cryo-EM or X-ray crystallography and highlight the potential issues surrounding them. MD has the power to help resolve these problems and indeed flexible fitting methods^39^, seem like a promising way forward to utilise this more directly at the model building stage. In the case of the pLGIC superfamily, the question of what truly represents the physiological open state has been subject to debate since 2015. Here, we report an MD-refined open and ion-conductive structure of the GlyR as a representative of this class. In light of our simulation results, and in revisiting the original cryo-EM density map together with the fitted atomic model 3JAE, we provide a new interpretation of the experimental data. We argue that the open state is stabilized by the L9’ residues, whose orientation is not resolved in the cryo-EM density map, being buried into hydrophobic pockets at the subunit interfaces, thus locking the receptor in an open state at the 9’ ring. We also suggest that the poor resolution of the cryo EM map at the P-2’ ring is the result of high dynamic flexibility in this region and the open state corresponds to an (on average symmetric) ensemble of possible asymmetric arrangements of the proline residues. Moreover, we argue that the -2’ constriction is *ca.* 1 Å smaller in radius than the fitted atomic model 3JAE suggests, which is underpinned by the original cryo-EM density in this region as well as our MD-simulation results and would be more in line with other open-like structures of the pLGIC superfamily. Since the non-polar 9’ position is highly conserved across the pLGICs (L or occasionally I/V), the new hydrophobic lock mechanism reported here is likely to play a key role in gating for the whole superfamily.

## Methods

### Homology Modelling

The human α1 GlyR model was constructed from the structure of *Danio rerio* α1 GlyR (glycine-bound open state; PDB: 3JAE^17^) as previously reported^16^. Briefly sequences were aligned using Muscle^40^ and adjusted manually. MODELLER version 9.16^41^ was used to generate 100 models and from of the intersection of the top 10 of both molecular PDF and DOPE score^42^, the model with the best QMEAN score^43^ was chosen for simulation. Hydrogens were added to the protein with the pdb2gmx tool of Gromacs 5.1^44^, with the default protonation state at pH = 7.4.

### Neurotransmitter Positioning and Parameterization

Glycine was positioned into all 5 orthosteric binding sites according to its position in the currently highest resolution glycine receptor structure; PDB:5TIN^19^, 2.6 Å resolution (note that glycine was bound to the receptor but no density was discernible in PDB:3JAE^17^, 3.9 Å resolution). We used the parameterization by Horn^45^ for zwitterionic amino acids with the AMBER99SB-ILDN force field. We also included a water molecule in the binding pocket which is resolved in PDB:5TIN and shown to be stable in MD-simulations^16,46^.

### System Set Up

Systems were set up using a coarse-grained approach with the Martini force field version 2.2 using martinize^47^ to coarse-grain the protein model and the “insane” programme^48^, using a hexagonal prism as the periodic simulation box. The protein was embedded in a 1-palmitoyl-2-oleoyl-sn-glycero-phosphocholine (POPC) bilayer, solvated in water and sodium and chloride ions were added to neutralise the net charge and simulate a physiological concentration of 150 mM. Gromacs 2018^44^ was used as molecular dynamics engine for all simulations. After energy minimisation, the system was equilibrated at 310 K and 1 bar for 10 ns. Simulation details were based on the recommended new-rf.mdp parameters^49^and appropriately adjusted. The final frame was then converted to all-atom resolution via the programme “backward”^50^. The back-mapped receptor was replaced by the original all-atom model with the glycine ligands and stable water molecules bound, resulting in a total of 136,334 atoms.

### All Atom Molecular Dynamics (MD) Simulations

#### Protocol

Each system was energy-minimised using the steepest descent algorithm until the maximum force fell below a tolerance of 1000 kJ mol^-1^ nm^-1^. The initial equilibration protocol is summarised in **Table 1.** The final frame after this initial equilibration period was used as the starting point for three independent simulations with different velocities, all of which exhibited hydrophobic collapse. To find a stable open channel structure, we extended the initial equilibration as follows. The ion channel pore was kept open via flat-bottom harmonic cross distance position restraints between the Cα atoms of pore-lining residues (pore restraints, see **Fig. 6**). The flat bottom potential only acts when the Cα atoms of opposite pore-lining residues come closer to each other than their distance in the open-state cryo-EM structure 3JAE with a force proportional to the difference in distance (5000 kJ mol^-1^ nm^-1^). This keeps the channel pore from collapsing initially and gives it enough freedom to find a stable configuration. This additional restrained simulation step with pore restraints was performed for 150 ns. The final frame was used as the starting configuration for three independent repeats, initiated with different velocities. All three repeats showed a stable open channel configuration. During both equilibration and production runs, flat-bottomed restraint terms were applied to the free glycine ligands in the binding site to prevent their dissociation, but still allow reasonable exploration of the pocket. These restraints act if the distance between Cα atoms of key residues in the binding site and the Cα atom of the glycine ligand exceed a threshold distance (namely, 10.60 Å to the (+)F207 Cα, 10.53 Å to the (-)F63 Cα and 9.15 Å to the (-)S129 Cα, with a force constant of 5000 kJ mol^-1^ nm^-1^).

**Table 1.** Initial equilibration protocol.

| Time | Ensemble | Position restraints on | Force constant<br>(kJ mol<sup>-1</sup> nm<sup>-1</sup>) |
| --- | --- | --- | --- |
| 100 ps | NVT | Protein heavy atoms, ligand heavy atoms,<br>crystal water oxygen | 1000 |
| 1 ns | NPT | Protein heavy atoms, ligand heavy atoms,<br>crystal water oxygen | 1000 |
| 10 ns | NPT | Protein backbone, ligand Cα atom, crystal water<br>oxygen | 1024 |
| 1 ns | NPT | Protein backbone, ligand Cα atom, crystal water<br>oxygen | 512 |
| 1 ns | NPT | Protein backbone, ligand Cα atom, crystal water<br>oxygen | 256 |
| 1 ns | NPT | Protein backbone, ligand Cα atom, crystal water<br>oxygen | 128 |
| 1 ns | NPT | Protein backbone, ligand Cα atom, crystal water<br>oxygen | 64 |

**Fig. 6.**
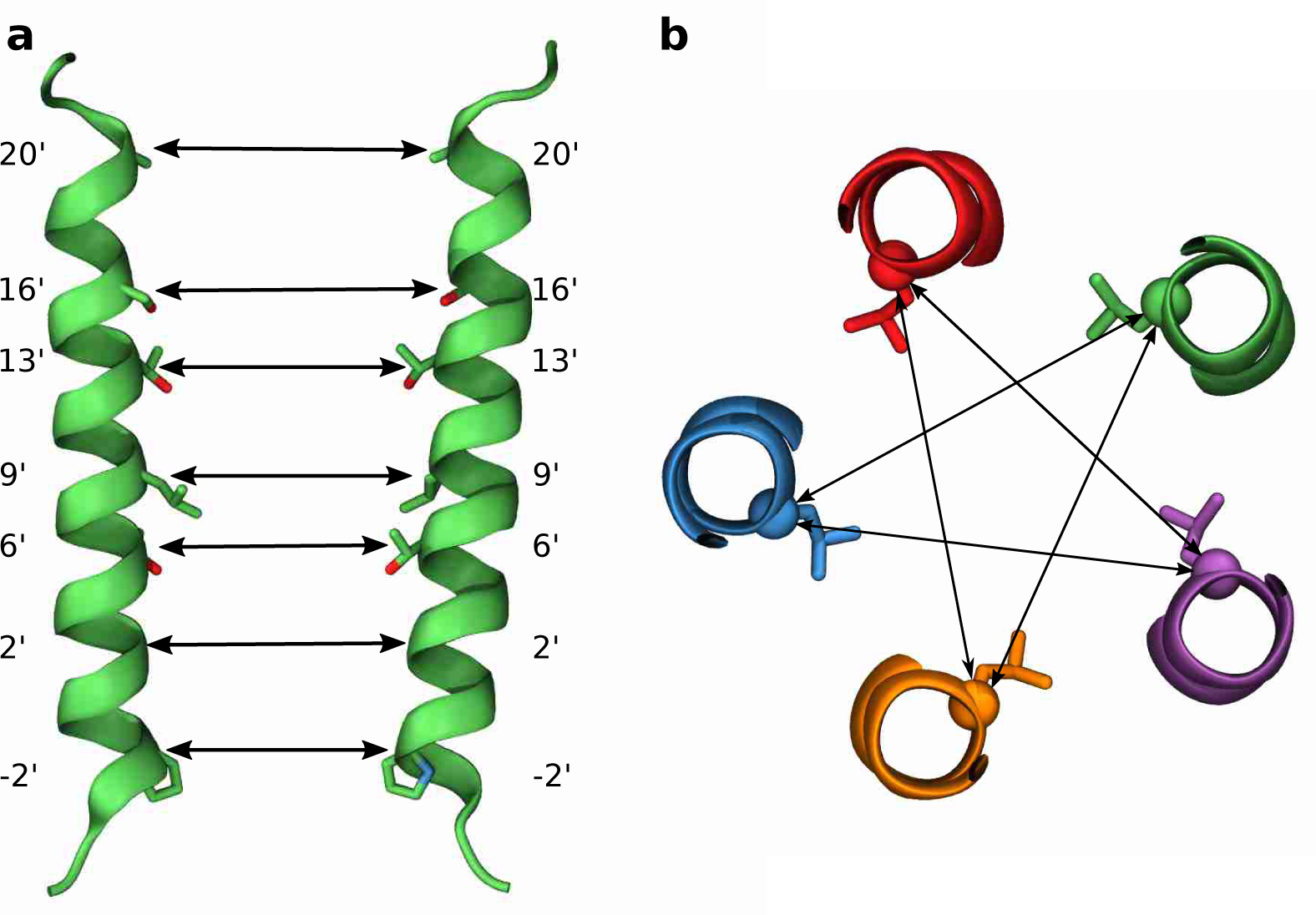
Pore restraints. **a** Two opposite pore lining M2 helices with seven pore lining residues. The double arrows indicate the pore cross distances on which the flat bottom pore restraints act. **b** Illustration of the pore cross distances at the L9’ gate (view from the extracellular towards the intracellular side). In total, there are five cross distances for the 9’ ring. The cross distances for the other pore lining residues are not shown for clarity, but similarly, there are five cross distances per pore lining residue position, so that in total 7 × 5 = 35 cross distances are restrained.

#### Simulation details

The force fields used were AMBER99SB-ILDN^51^ for protein and ions, TIP3P for water^52^, Slipids^53-55^ for lipids and the parameters for zwitterionic amino acids by Horn^45^ for the glycine ligands. Periodic boundary conditions in all three spatial dimensions were applied to mimic bulk conditions. The Bussi thermostat^56^ was used to maintain the temperature constant at physiological 310 K using a coupling constant of 0.1ps. In all simulations with constant pressure, a barostat was coupled to the system in a semi-isotropic manner to maintain a pressure of 1 bar. The time constant used for pressure coupling was 1 ps and the isothermal compressibility set to 4.5 × 10-5 bar-1. For the initial equilibration, the Berendsen barostat^57^ was used and the Parrinello-Rahmann barostat^58^ for production runs as well as the simulation period with pore restraints. The van der Waals interactions were cut off at 10 Å and a dispersion correction was applied to energy and pressure. Electrostatic interactions were treated using the smooth Particle Mesh Ewald (PME) method^59,60^, where the real space contribution was cut off at 10 Å and the reciprocal energy term obtained in k-space was calculated on a grid with 1.2 Å spacing using 4th order B-splines for interpolation. The Verlet cut-off scheme was used to generate a pair-list with buffering using a tolerance of 0.005 kJ/mol/ps for pair-interactions per particle. All bonds involving hydrogens were constrained with the LINCS algorithm^61^, allowing for a time step of 2 fs (except in the first dynamics run (NVT-equilibration), where a shorter time step of 1 fs was used). Coordinates were written out every 10 ps.

### Analysis and Visualization

Pore profiles, volumes and hydrophobicity were calculated with CHAP^62^. Hydrophobicity was calculated according to the Wimley and White scale^63,64^. Trajectory analysis was performed with MDAnalysis^65^ and in-house scripts. The local resolution of the cryo-EM density map was estimated with RESMAP^66^, the real space correlation between atomic model and map was calculated with Phenix^67^. Visualization was performed in VMD^68^ and UCSF Chimera^69^.

## Acknowledgements

We thank our colleagues for useful discussions and analysis scripts, especially Hannah Smith, Gianni Klesse and Shanlin Rao. MAD received funding by the Oxford Wolfson Marriott Graduate Scholarship in Biochemistry, the Studienstiftung des deutschen Volkes, the Goodger and Schorstein Research Scholarship in Medical Sciences as well as the Vice-Chancellors’ fund. This project made use of computation time on JADE (EP/P020275/1) via HECBioSim (http://www.hecbiosim.ac.uk), supported by EPSRC (grant no. EP/R029407/1). PCB thanks the MRC for funding (BB/S001247/1).

## Author Contributions

PCB and MAD conceived the study. MAD designed the simulation protocol, performed and analysed the MD simulations as well as the analysis of the cryo-EM map and atomic model 3JAE. MAD and PCB interpreted the results and wrote the paper.

## Competing Interests

The authors declare no competing interests.

## Materials and Correspondence

A zipped archive file is included as **Supplementary Data** that contains the trajectory referred to as repeat 1 in the main manuscript with frames written out every 1ns, the first frame of the MD trajectory in GROMACS format and a pdb file of the representative open structure. All requests for materials and correspondence should be addressed to Philip Biggin.

## References

1 Jensen, A. A., Frølund, B., Liljefors, T. & Krogsgaard-Larsen, P. Neuronal nicotinic acetylcholine receptors: Structural revelations, target identifications and therapeutic inspirations. J. Med. Chem. 48, 4705–4745, (2005).

2 Gotti, C., Zoli, M. & Clementi, F. Brain nicotinic acetylcholine receptors: native subtypes and their relevance. Trends Pharmacol. Sci. 27, 482–491, (2006).

3 Van Arnam, E. B. & Dougherty, D. A. Functional probes of drug-receptor interactions implicated by structural studies: Cys-loop receptors provide a fertile testing ground. J. Med. Chem. 57, 6289–6300, (2014).

4 Nemecz, Á., Prevost, Marie S., Menny, A. & Corringer, P.-J. Emerging molecular mechanisms of signal transduction in pentameric ligand-gated ion channels. Neuron 90, 452–470, (2016).

5 Katz, B. & Thesleff, S. A study of the densitization produced by acetylcholine at the motor end-plate. J. Physiol. 138, 63–80, (1957).

6 Cecchini, M. & Changeux, J.-P. The nicotinic acetylcholine receptor and its prokaryotic homologues: Structure, conformational transitions & allosteric modulation. Neuropharmacology 96, Part B, 137–149, (2015).

7 Nys, M., Kesters, D. & Ulens, C. Structural insights into Cys-loop receptor function and ligand recognition. Biochemical Pharmacology 86, 1042–1053, (2013).

8 Burgos, C. F., Yévenes, G. E. & Aguayo, L. G. Structure and Pharmacologic Modulation of Inhibitory Glycine Receptors. Mol. Pharmacol. 90, 318, (2016).

9 Lynch, J. W. Molecular structure and function of the glycine receptor chloride channel. Physiol. Rev. 84, 1051–1095, (2004).

10 Lynch, J. W. Native glycine receptor subtypes and their physiological roles. Neuropharmacology 56, 303–309, (2009).

11 Lynch, J. W. & Callister, R. J. Glycine receptors: a new therapeutic target in pain pathways. Curr. Opin. Investig. Drugs 7, 48–53, (2006).

12 Bode, A. & Lynch, J. The impact of human hyperekplexia mutations on glycine receptor structure and function. Mol. Brain 7, 2, (2014).

13 Lynch, J. W., Zhang, Y., Talwar, S. & Estrada-Mondragon, A. in Adv. Pharmacol. Vol. 79 (eds Dominic P. Geraghty & Lachlan D. Rash) 225–253 (Academic Press, 2017).

14 Lape, R., Colquhoun, D. & Sivilotti, L. G. On the nature of partial agonism in the nicotinic receptor superfamily. Nature 454, 722–727, (2008).

15 Lape, R., Plested, A. J. R., Moroni, M., Colquhoun, D. & Sivilotti, L. G. The α1K276E startle disease mutation reveals multiple intermediate states in the gating of glycine receptors. J. Neurosci. 32, 1336–1352, (2012).

16 Safar, F. et al. The startle disease mutation E103K impairs activation of human homomeric α1 glycine receptors by disrupting an intersubunit salt bridge across the agonist binding site. J. Biol. Chem. 292, 5031–5042, (2017).

17 Du, J., Lu, W., Wu, S., Cheng, Y. & Gouaux, E. Glycine receptor mechanism elucidated by electron cryo-microscopy. Nature 526, 224–229, (2015).

18 Huang, X., Chen, H. & Shaffer, P. L. Crystal structures of human GlyRα3 bound to ivermectin. Structure 25, 945–950.e942, (2017).

19 Huang, X. et al. Crystal structures of human glycine receptor α3 bound to a novel class of analgesic potentiators. Nat. Struct. Mol. Biol. 24, 108–113, (2017).

20 Huang, X., Chen, H., Michelsen, K., Schneider, S. & Shaffer, P. L. Crystal structure of human glycine receptor alpha3 bound to antagonist strychnine. Nature 526, 277–280, (2015).

21 Thompson, A. J., Lester, H. A. & Lummis, S. C. R. The structural basis of function in Cys-loop receptors. Quart. Rev. Biophys. 43, 449–499, (2010).

22 Sauguet, L. et al. Structural basis for ion permeation mechanism in pentameric ligand-gated ion channels. The EMBO J. 32, 728, (2013).

23 Hibbs, R. & Gouaux, E. Principles of activation and permeation in an anion-selective Cys-loop receptor. Nature 474, 54–60, (2011).

24 Gonzalez-Gutierrez, G., Wang, Y., Cymes, G. D., Tajkhorshid, E. & Grosman, C. Chasing the open-state structure of pentameric ligand-gated ion channels. J. Gen. Physiol. 149, 1119, (2017).

25 Beckstein, O., Biggin, P. C. & Sansom, M. S. P. A hydrophobic gating mechanism for nanopores. J. Phys. Chem. B. 105, 12902–12905, (2001).

26 Beckstein, O. & Sansom, M. S. P. A hydrophobic gate in an ion channel: the closed state of the nicotinic acetylcholine receptor. Phys. Biol. 3, 147–159, (2006).

27 Aryal, P., Sansom, M. S. P. & Tucker, S. J. Hydrophobic gating in ion channels. J. Mol. Biol. 427, 121–130, (2015).

28 Trick, J. L. et al. Functional Annotation of Ion Channel Structures by Molecular Simulation. Structure (London, England : 1993) 24, 2207–2216, (2016).

29 Cerdan, A. H., Martin, N. É. & Cecchini, M. An ion-permeable state of the glycine receptor captured by molecular dynamics. Structure 26, 1555–1562, (2018).

30 Cheng, M. H. & Coalson, R. D. Energetics and Ion permeation characteristics in a glutamate-gated chloride (GluCl) receptor channel. J. Phys. Chem. B 116, 13637–13643, (2012).

31 Calimet, N. et al. A gating mechanism of pentameric ligand-gated ion channels. Proceedings of the National Academy of Sciences, (2013).

32 Yoluk, O., Lindahl, E. & Andersson, M. Conformational gating dynamics in the GluCl anion-selective chloride channel. ACS Chem. Neurosci. 6, 1459–1467, (2015).

33 Filatov, G. N. & White, M. M. The role of conserved leucines in the M2 domain of the acetylcholine-receptor in channel gating. Mol. Pharmacol. 48, 379–384, (1995).

34 Labarca, C. et al. Channel gating governed symmetrically by conserved leucine residues in the M2 domain of nicotinic receptors. Nature 376, 514–516, (1995).

35 Beckstein, O. & Sansom, M. S. P. The influence of geometry, surface character, and flexibility on the permeation of ions and water through biological pores. Phys. Biol. 1, 42–52, (2004).

36 Keramidas, A., Moorhouse, A. J., Pierce, K. D., Schofield, P. R. & Barry, P. H. Cation-selective Mutations in the M2 Domain of the Inhibitory Glycine Receptor Channel Reveal Determinants of Ion-Charge Selectivity. J. Gen. Physiol. 119, 393, (2002).

37 Basak, S., Gicheru, Y., Rao, S., Sansom, M. S. P. & Chakrapani, S. Cryo-EM reveals two distinct serotonin-bound conformations of full-length 5-HT3A receptor. Nature 563, 270–274, (2018).

38 Polovinkin, L. et al. Conformational transitions of the serotonin 5-HT3 receptor. Nature 563, 275–279, (2018).

39 Igaev, M., Kutzner, C., Bock, L. V., Vaiana, A. C. & Grubmüller, H. Automated cryo-EM structure refinement using correlation-driven molecular dynamics. eLife 8, e43542, (2019).

40 Edgar, R. C. MUSCLE: Multiple sequence alignment with high accuracy and high throughput. Nucleic Acids Research 32, 1792–1797, (2004).

41 Webb, B. & Sali, A. Comparative Protein Structure Modeling Using Modeller. Curr. Prot. Bioinf. 5, 5.61–65.66.32, (2014).

42 Shen, M. Y. & Sali, A. Statistical potential for assessment and prediction of protein structures. Protein Sci 15, 2507–2524, (2006).

43 Benkert, P., Kunzli, M. & Schwede, T. QMEAN server for protein model quality estimation. Nucleic Acids Res, (2009).

44 Abraham, M. J. et al. GROMACS: High performance molecular simulations through multi-level parallelism from laptops to supercomputers. SoftwareX 1–2, 19–25, (2015).

45 Horn, A. H. C. A consistent force field parameter set for zwitterionic amino acid residues. J. Mol. Mod. 20, 2478, (2014).

46 Yu, R. et al. Agonist and antagonist binding in human glycine receptors. Biochemistry 53, 6041–6051, (2014).

47 de Jong, D. H. et al. Improved parameters for the martini coarse-grained protein force field. J. Chem. Theory Comput. 9, 687–697, (2013).

48 Wassenaar, T. A., Ingólfsson, H. I., Böckmann, R. A., Tieleman, D. P. & Marrink, S. J. Computational lipidomics with insane: A versatile tool for generating custom membranes for molecular simulations. J. Chem. Theor. Comput. 11, 2144–2155, (2015).

49 de Jong, D. H., Baoukina, S., Ingólfsson, H. I. & Marrink, S. J. Martini straight: Boosting performance using a shorter cutoff and GPUs. Comp. Phys. Comms. 199, 1–7, (2016).

50 Wassenaar, T. A., Pluhackova, K., Böckmann, R. A., Marrink, S. J. & Tieleman, D. P. Going backward: A flexible geometric approach to reverse transformation from coarse grained to atomistic models. J. Chem. Theor. Comput. 10, 676–690, (2014).

51 Lindorff-Larsen, K. et al. Improved side-chain torsion potentials for the Amber ff99SB protein force field. Proteins: Struc. Func. Genet. 78, 1950–1958, (2010).

52 Jorgensen, W. L., Chandresekhar, J., Madura, J. D., Impey, R. W. & Klein, M. L. Comparison of simple potential functions for simulating liquid water. J. Chem. Phys. 79, 926–935, (1983).

53 Jämbeck, J. P. M. & Lyubartsev, A. P. Another piece of the membrane puzzle: Extending Slipids further. J. Chem.Theory Comp. 9, 774–784, (2013).

54 Jämbeck, J. P. M. & Lyubartsev, A. P. An extension and further validation of an all-atomistic force field for biological membranes. J. Chem.Theory Comp. 8, 2938–2948, (2012).

55 Jämbeck, J. P. M. & Lyubartsev, A. P. Derivation and systematic validation of a refined all-atom force field for phosphatidylcholine lipids. J. Phys. Chem. B 116, 3164–3179, (2012).

56 Bussi, G., Donadio, D. & Parrinello, M. Canonical sampling through velocity rescaling. J. Chem. Phys. 126, (2007).

57 Berendsen, H. J. C., Postma, J. P. M., van Gunsteren, W. F., DiNola, A. & Haak, J. R. Molecular dynamics with coupling to an external bath. J. Chem. Phys. 81, 3684–3690, (1984).

58 Parinello, M. & Rahman, A. Polymorphic transitions in single crystals - a new molecular dynamics method. J. Appl. Phys. 52, 7182–7190, (1981).

59 Essman, U. et al. A smooth particle mesh Ewald method. J. Chem. Phys. 103, 8577–8593, (1995).

60 Darden, T., York, D. & Pedersen, L. Particle mesh Ewald - an N.log(N) method for Ewald sums in large systems. J. Chem. Phys. 98, 10089–10092, (1993).

61 Hess, B., Bekker, J., Berendsen, H. J. C. & Fraaije, J. G. E. M. LINCS: A linear constraint solver for molecular simulations. J. Comp. Chem. 18, 1463–1472, (1997).

62 Klesse, G., Rao, S., Sansom, M. S. P. & Tucker, S. J. CHAP: a versatile tool for the structural and functional annotation of ion channel pores. bioRxiv, 527275, (2019).

63 Wimley, W. C., Creamer, T. P. & White, S. H. Solvation energies of amino acid side chains and backbone in a family of host-guest pentapeptides. Biochemistry 35, 5109–5124, (1996).

64 Wimley, C. W. & White, S. H. Experimentally determined hydrophobicity scale for proteins at membrane interfaces. Nature Struct. Biol. 3, 842–848, (1996).

65 Michaud-Agrawal, N., Denning, E. J., Woolf, T. & Beckstein, O. MDAnalysis: A toolkit for the analysis of molecular dynamics simulations. J. Comput. Chem. 32, 2319–2327, (2011).

66 Kucukelbir, A., Sigworth, F. J. & Tagare, H. D. Quantifying the local resolution of cryo-EM density maps. Nat. Methods 11, 63–65, (2014).

67 Adams, P. D. et al. PHENIX: a comprehensive Python-based system for macromolecular structure solution. Acta Crystallographica Section D 66, 213–221, (2010).

68 Humphrey, W., Dalke, A. & Schulten, K. VMD - Visual molecular dynamics. J. Mol. Graph. 14, 33–38, (1996).

69 Pettersen, E. F. et al. USCF chimera - A visualization system for exploratory research and analysis. J. Comp. Chem. 25, 1605–1612, (2004).

