## Supplementary Figures for "A Refined Open State of the Glycine Receptor Obtained Via Molecular Dynamics Simulations"

*To whom correspondence should be addressed.

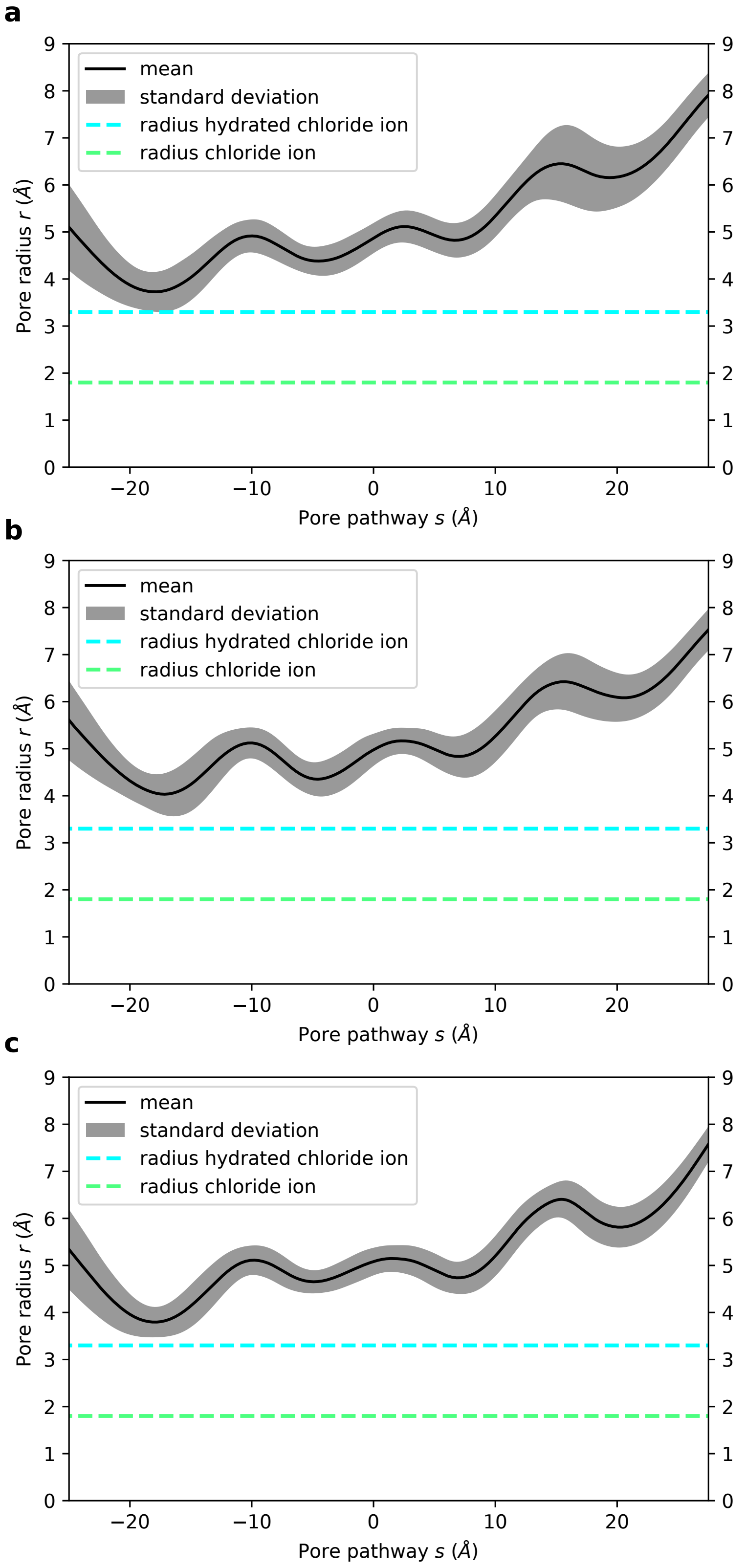


**Supplementary Figure 1 | Pore profiles from three independent simulations with a stable open state.** Time averaged pore radius profiles of the transmembrane domain obtained from three independent simulations (of 300 ns length each) initiated with different velocities from the system obtained after 150 ns equilibration with pore restraints. No restraints were applied to keep the pore open. The radii of a dehydrated and a hydrated chloride ion are indicated by dashed green and cyan lines, respectively. The pore is positioned in this and subsequent figures such that the L9' ring is located at s = 0. In all three repeats, the structure from the simulation is physically open and theoretically allows permeation of hydrated chloride ions.

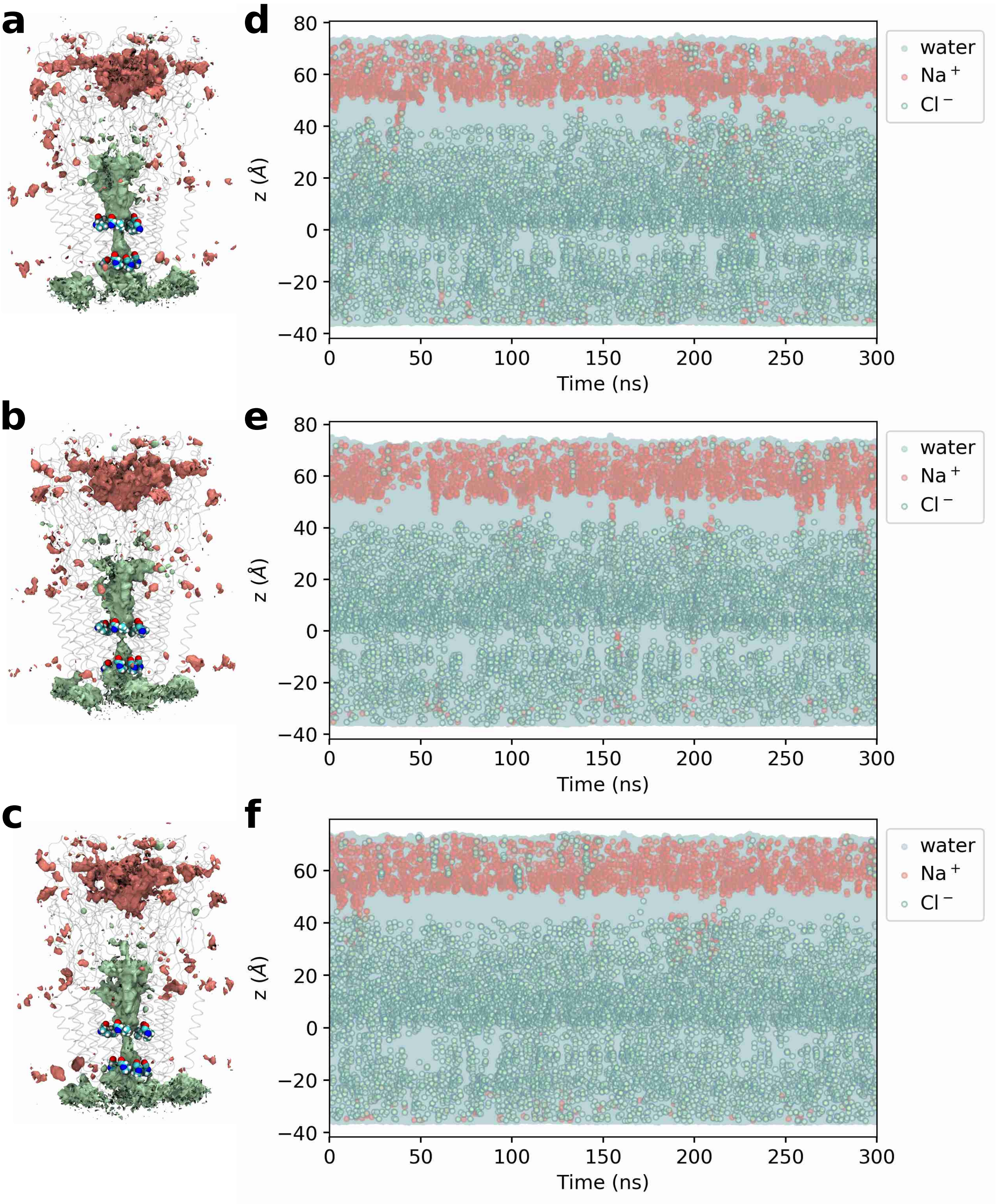


**Supplementary Figure 2 | Ion densities and trajectories of water and ions within 5 Å of the channel axis for simulations with a stable open state.** (a-c) Densities of chloride (green) and sodium (red) ions over 300 ns. The isosurfaces represent a density value of 0.5 particles/nm$^3$. Transparent ribbons indicate the receptor backbone. The L9' ring and P-2' ring residues are shown in van der Waals representation. The densities prove chloride occupancy in the transmembrane pore and, moreover, show selectivity for chloride over sodium in this region. (d-f) Trajectories of water, chloride and sodium ion z-coordinates within 5 Å of the channel axis inside the pore over 300 ns (represented by blue, green and red circles, respectively). The 0 point of the z-axis is positioned at the L9' ring and the receptor structures in the left panels are aligned and scaled correspondingly to allow for spatial orientation along the z-axis. The whole channel pore is wetted throughout the simulations (dewetted regions would appear as white stretches). While chloride frequently penetrates the transmembrane pore, sodium only does so very rarely, again demonstrating the chloride selectivity for this channel.

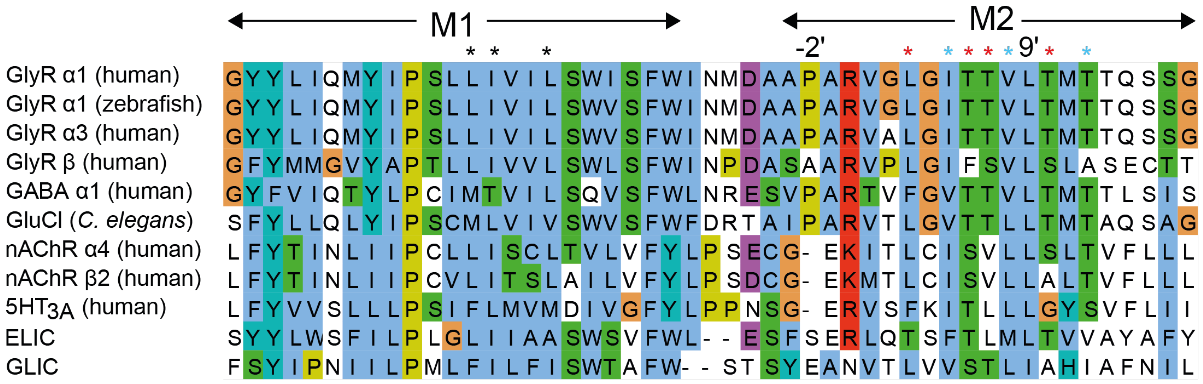


**Supplementary Figure 3 | Multiple sequence alignment of pLGIC representatives.** The conservation of the leucine at L9' is well described. Less well appreciated is the functional conservation of the residues that form the hydrophobic pocket for the leucine in the open state formed by I257, V260 and T264 (I5', V8' and T12' respectively and shown as cyan asterisks) belonging to the M2 helix of the principal (+) subunit, L255, T258, T259, T262 (L3', T6', T7' and T10' respectively and shown as red asterisks) belonging to the M2 helix of the complementary (-) subunit as well as L233, I234 and L237 of the M1 helix of the complementary (-) subunit, shown as black asterisks.

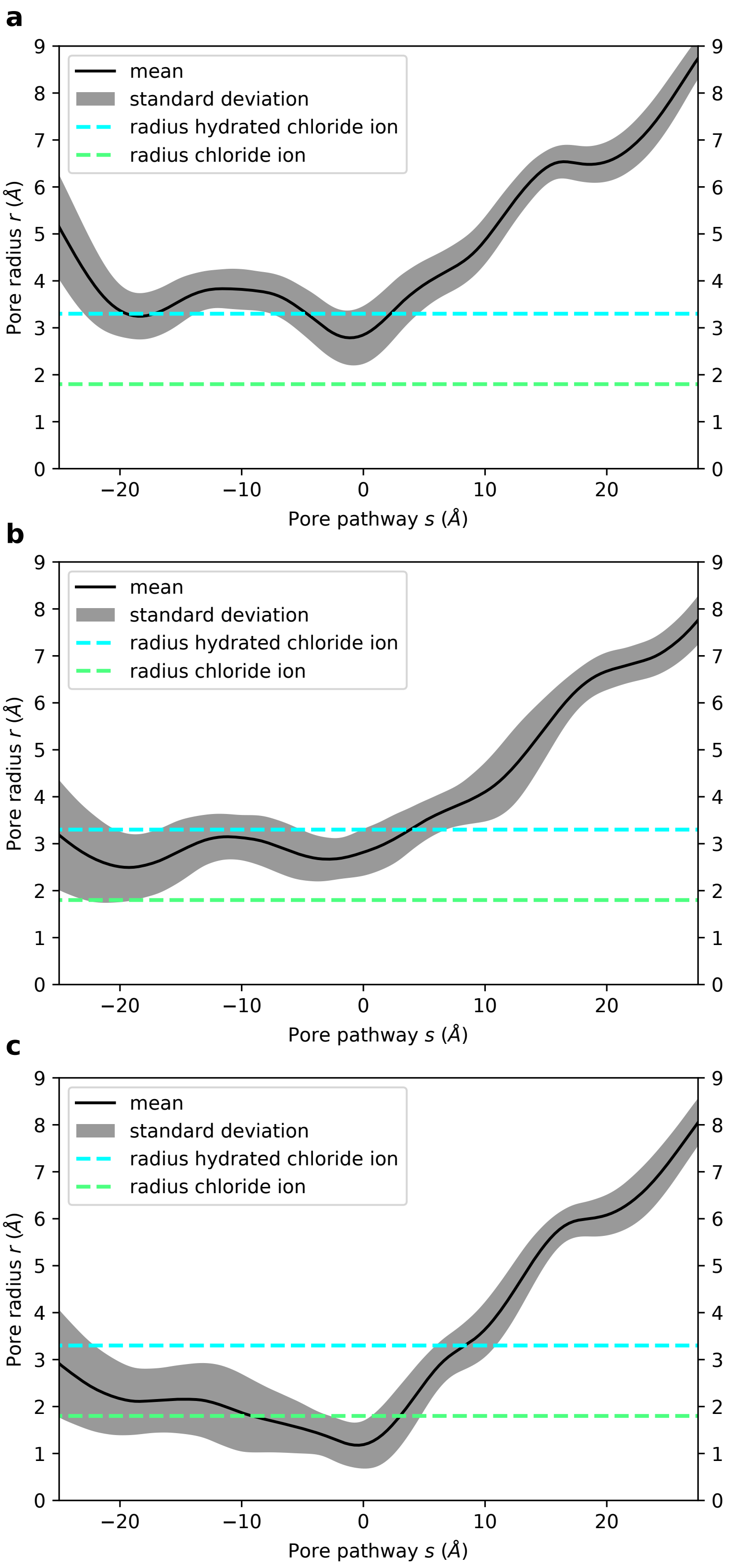


**Supplementary Figure 4 | Pore profiles from three independent simulations with a collapsed state.** Time averaged pore radius profiles of the transmembrane domain obtained from three independent simulations (of 300 ns length each) initiated with different velocities from the system obtained immediately after the short initial equilibration before the structure is relaxed via the application of minimally invasive pore restraints. The radii of a dehydrated and a hydrated chloride ion are indicated by dashed green and cyan lines, respectively. The L9’ and P-2’ rings are the main constriction points. If at all, only partially dehydrated chloride ions could theoretically permeate through the pore.

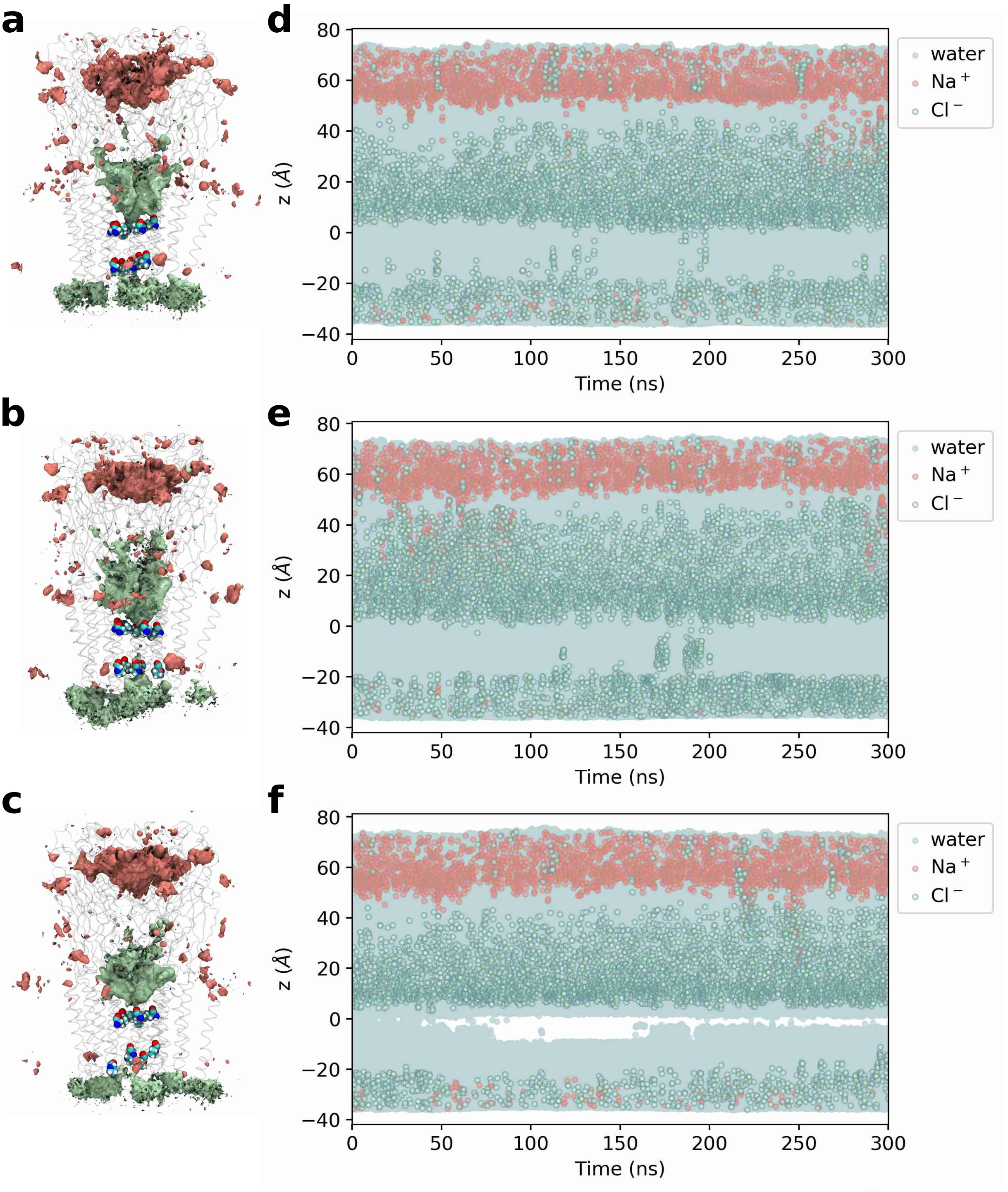


**Supplementary Figure 5 | Ion densities and trajectories of water and ions within 5 Å of channel axis for simulations with a collapsed state.** (a-c) Densities of chloride (green) and sodium (red) ions over 300 ns production run. The isosurfaces shown represent a density value of 0.5 particles/nm$^3$. Transparent ribbons indicate the receptor backbone, The L9' ring and P-2' ring residues are shown in van der Waals representation. The densities show chloride occupancy in the transmembrane pore and moreover, show selectivity for chloride over sodium in this region. (d-f) Trajectories of water as well as chloride and sodium ion z-coordinates within 5 Å of the channel axis inside the pore over 300 ns (represented by blue, green and red circles, respectively). The 0 point of the z-axis is positioned at the L9' ring and the receptor structures in the left panels are aligned and scaled correspondingly to allow for spatial orientation along the z-axis. The white stretches in (c) demonstrate local dewetting of the pore. Chloride ions access the transmembrane pore much less frequently than in the stable open state simulations.

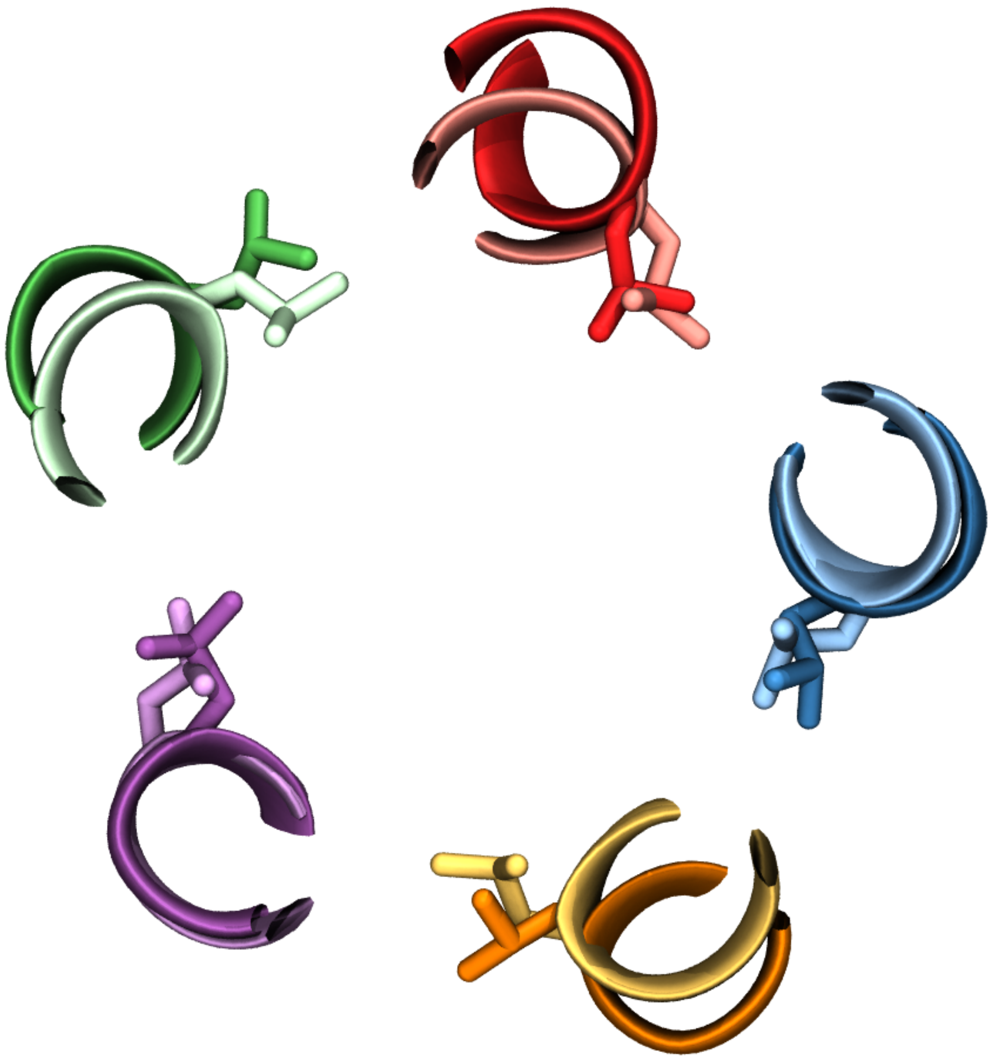

**Supplementary Figure 6 | Comparison of the leucine gate between the open state glycine and the 5HT$_3$ receptor.** The open state model (dark colours) was fitted to the C$\alpha$ atoms of L9' of the 5HT$_3$ structure (light colours and PDB code: 6HIN). Shown is a top down view of a horizontal slice at the L9' position and highlights the similarity in position of the leucines at this position in an open state.

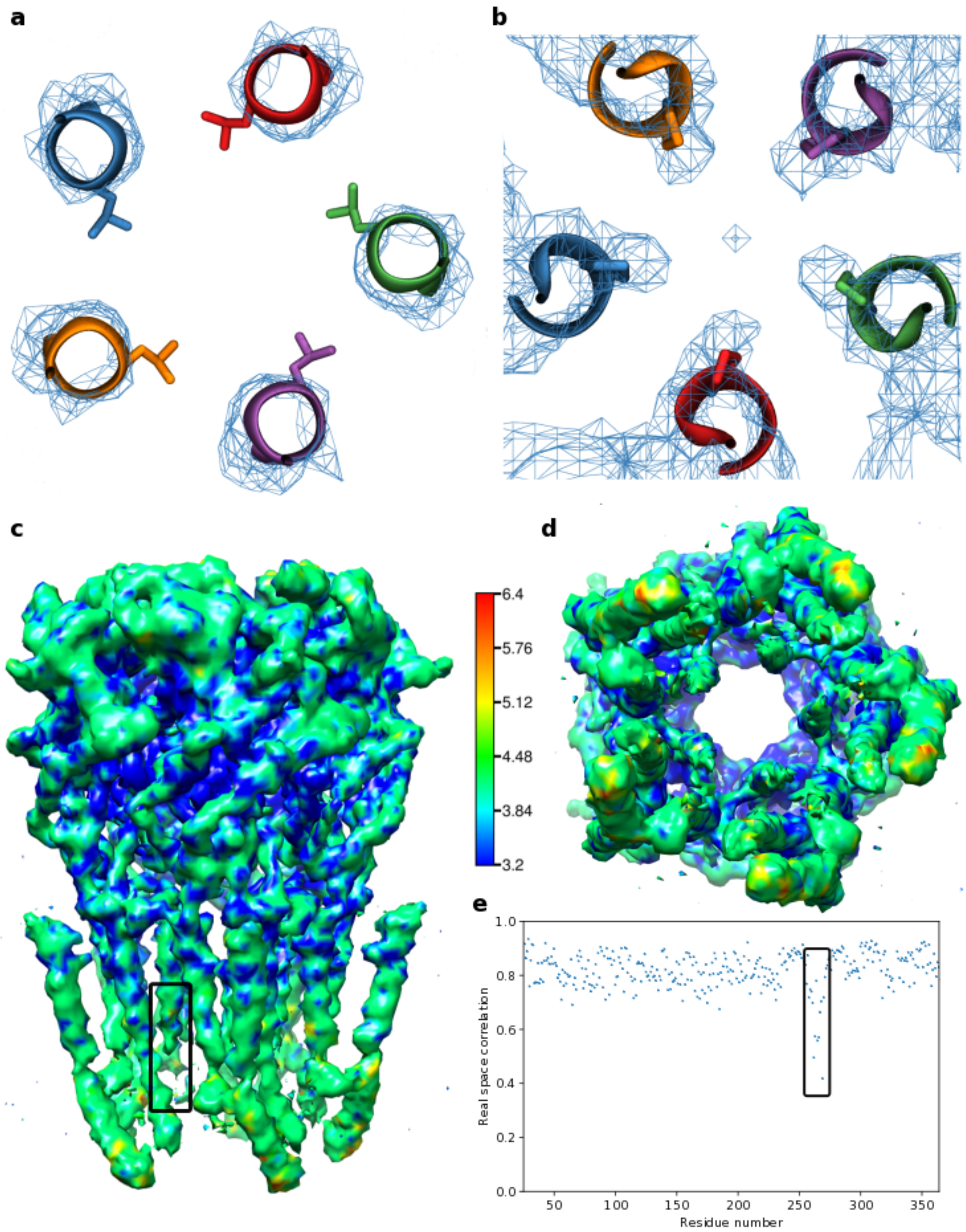


**Supplementary Figure 7 | Cryo-EM density map and fitted model of 3JAE structure.** (a) Electron density of the L9' gate region, contoured at an isosurface with a value of 6.0 together with the fitted model structure for 3JAE (viewed from the extracellular towards the intracellular side). The density is insufficient to indicate the leucine side chain orientation and is therefore purely a result of computational modelling. (b) Electron density of the P-2' gate region, contoured at an isosurface with a value of 5.8 together with the fitted model structure for 3JAE (viewed from intracellular towards the extracellular side). The density suggests a narrower –2' pore restriction as the fitted model. (c and d) Electron density map coloured according to local resolution estimated using RESMAP and visualized with Chimera (side view (c) and view from the intracellular side towards the extracellular side (d)). The local resolution is particularly poor in the lower pore lining M2 helix region and its linker to the M1 helix (highlighted by the box) which suggests a higher dynamic flexibility in this region. (e) Real-space correlation between the fitted atomic model and the electron density map calculated with PHENIX. The correlation is particularly poor in the lower half of the pore lining M2 helix and its linker to the M1 helix (highlighted by black box), indicating that the quality of the atomic model is particularly poor in this region.

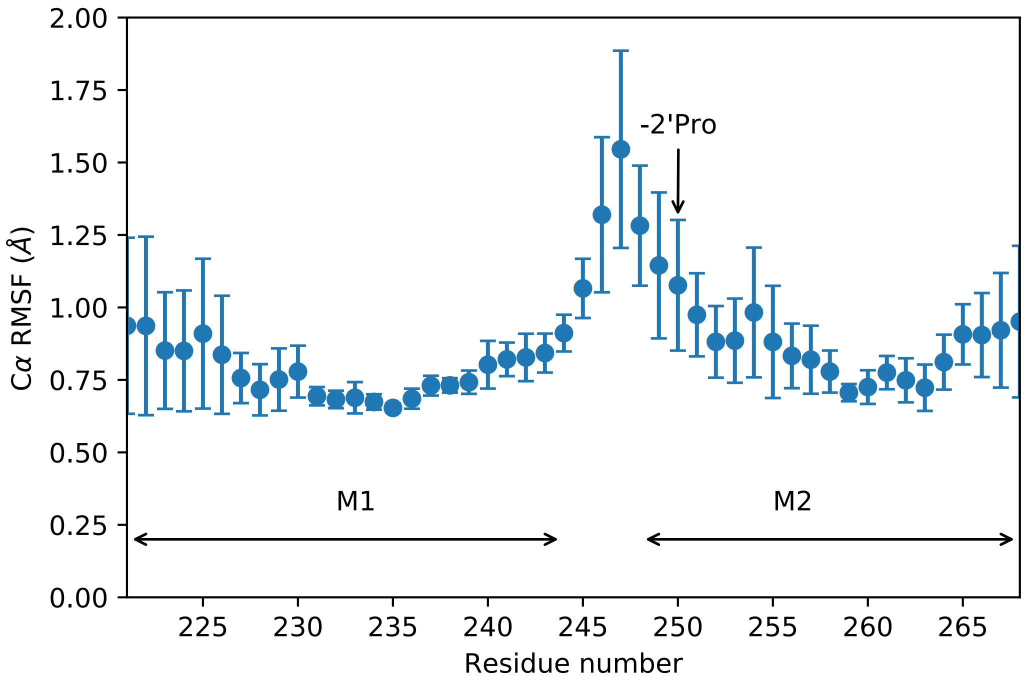


**Supplementary Figure 8 | Root mean square fluctuation (RMSF) of Cα atoms of M1 and M2 helices.** RMSF of Cα atoms of the M1 and M2 helices averaged over all five subunits with standard deviation as error bars. The values corresponding to the region near the intracellular pore opening of the pore lining M2 helix (where the P-2' gate is located) are very high compared to typical values for alpha-helical Cα atoms of around 0.7 Å. This indicates a high conformational flexibility that is accessible at a physiological temperature of 37 °C and can explain the poor electron density in the cryo-EM map in this region.
